# NOMA: a high-throughput microchip for robust, sequential measurements of secretions from the same single-cells

**DOI:** 10.1101/2021.01.21.427049

**Authors:** Meimei Liu, Yahui Ji, Fengjiao Zhu, Xue Bai, Linmei Li, Hua Xie, Xianming Liu, Yong Luo, Tingjiao Liu, Bingcheng Lin, Yao Lu

## Abstract

Despite recent advances in single-cell analysis technologies, lacking simple methods to keep the live single-cells traceable for longitudinal detection reliably poses a significant obstacle in single-cell secretion analysis. Here we reported the high-density NOMA (narrow-opening microwell array) microchip that realized the retention of **≥**97% of trapped single cells in dedicated spatial locations during repetitive detection procedures, verified with both adherent and suspension cells by two researchers independently. We applied it in monitoring single-cell protein secretions sequentially from the same single cells, and we found the digital protein secretion patterns dominate the protein secretion. We also demonstrated the microchip for longitudinally tracking IL-8 and the CD81+EV secretions from the same single-cells over days, which revealed the presence of “super secretors” within the cell population be more persistent to secrete protein or extracellular vesicle for an extended period. The NOMA platform reported here is simple, robust, and easy to operate for tracking sequential measurements from the same single cells, representing a novel and informative tool to inspire new observations in biomedical research.

**Table of Contents Graphic:** 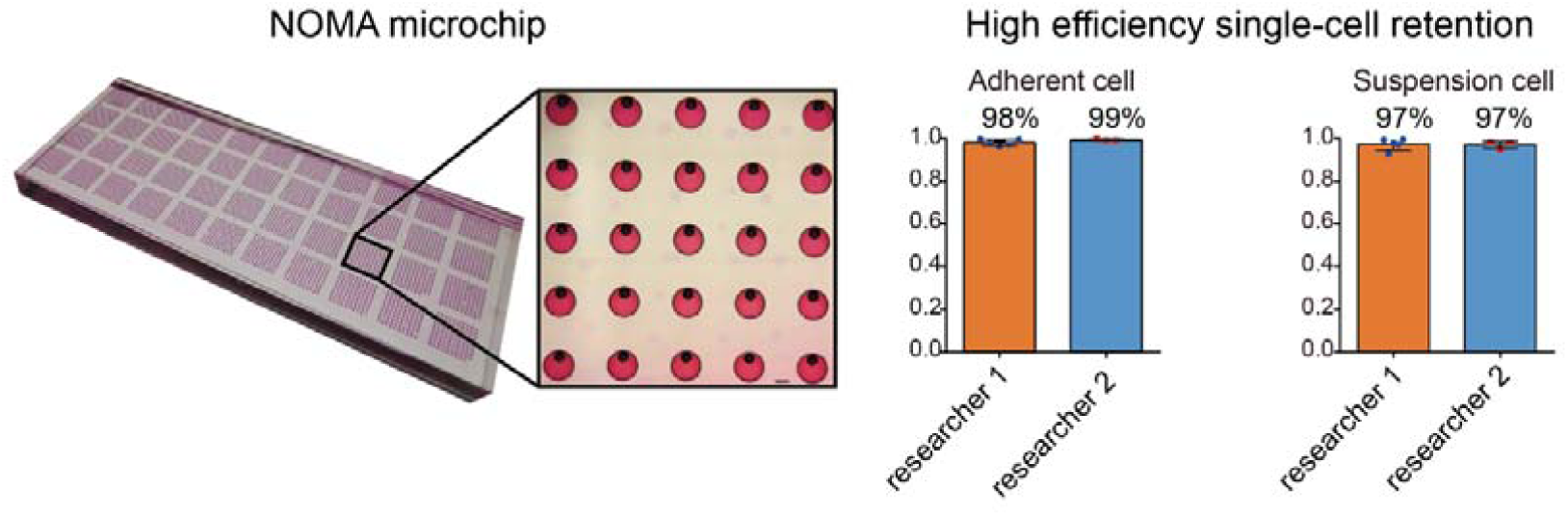

## Introduction

Cell heterogeneity plays a critical role in shaping cell physiology and fates in most biological processes(*1, 2*). Different single-cell analysis technologies, including genomics(*3*), transcriptomics(*4–6*), and proteomics(*7–9*), etc.(*10*), are rapidly developing to decode cell heterogeneity at different levels. As a critical player in mediating intercellular communications, cell secretion also exhibits high heterogeneity(*11*). For example, the polyfunctional immune cells, which could secrete multiple cytokines simultaneously, can be a highly predictive indicator of protective immunity or patient responses in immunotherapy clinical trials(*11–13*). Discriminating the secretory phenotypes at the single-cell resolution provides a direct way to evaluate the heterogeneity in cell states and activation under different physiological and diseased conditions, which could be of critical importance in basic cell biology, biomarker discovery, and disease diagnosis (*14–21*). Many efforts have been made to improve the multiplexity in secretion profiling at a single-cell resolution(*18, 22–24*). For example, Wang et al. introduced a multiplexed in situ tagging (MIST) strategy that combines ssDNA encoded microbead arrays with a DNA encoded antibody library to realize tens of protein profiling in single cells through multicolor and multicycle successive imaging(*23*). By combining spatial and spectral encoding, Lu et al. realized the 42 multiplexed protein secretion profiling from individual macrophage cells, representing the highest multiplexed capability to date for secretomic detection in single cells(*18*).

Besides high multiplexity, dynamic change is another inherent nature of cell secretion (*14, 25–27*), bringing more complexity in detection. Resolving the secretion information from the same single-cells at different time points could provide a new dimension of information for an in-depth understanding of how individual cells would evolve upon external stimulation, attracting increasing attention in recent years(*25, 28–31*). Imaging-based methods, including SPR(*32, 33*), scattered light(*30*), photonic crystal resonant imaging(*29*), optofluidic(*34*), etc., have been developed to visualize the secretion information continuously. Due to the limited binding sites and detection regions, these methods usually suffer from low throughput (several), low multiplexity, or a short detection time (within hours). To obtain multiplexed secretion information from more single-cells, high-throughput single-cell analysis platforms need to be employed. Currently used high throughput single-cell secretion analysis tools, including ELISpot (Fluorospot)(*35*), intracellular cytokine staining (ICS) flow cytometry with either fluorescence or mass spectrometry (CyTOF) detection(*7*), can only be used in snapshot measurements at an endpoint due to that they can not retain live cells after the detection. Microchip-based platforms, such as microengraving(*14, 31, 36, 37*), single-cell barcode microchips (*16, 24, 38*), can track single-cell secretions at different time points due to the cell adherence on the PDMS microchips. However, these methods suffered from significant cell loss during repetitive detection steps. It is reported that around 43% of single cells would be lost during sequential measurements, even for adherent cells, which brings the cell loss problem and the potential cell bias in the process. Junkin et al. reported a microvalve-based integrated microfluidic device to profile single-cell secretion dynamics (*25*). However, due to the complexity of microvalve control, the number of single cells profiled simultaneously is relatively low (only dozens), and the operation procedures are very complicated, making it difficult for widespread use. DropMap employed the high throughput droplets to classify single cells into stationary two-dimensional droplets array, enabling sensitive kinetic secretion analysis(*26*). But the limited oxygen, nutrient supply in the pL-droplets limited the cell culture within 12 h. Besides, the closed design makes it challenging to introduce perturbation during the process. Despite these recent advances, it remains an unmet need to reliably keep the live single-cells traceable for longitudinal detection in a simple way. Minimizing cell loss during sequential readout with the same single-cells is the most critical and technically challenging.

To address the need, herein, we developed the high-density NOMA microchip to realize the retaining of ≥97% of trapped single cells to keep single-cells traceable in dedicated spatial positions during repetitive secretion detection procedures. The high retention efficiency was verified with both adherent and suspension cells by two researchers independently, demonstrating the high fidelity of the microchip performance. The platform makes it possible to track the information from the same live single-cells sequentially. We demonstrated the potential of the NOMA microchip in evaluating secretion functions from the same single-cells by two applications: measuring the protein secretion sequentially from the same single cells, and longitudinally tracking of the single-cell integrative secretions over days.

## Experimental methods

### Cell culture

SCC25 (ATCC(American Type Culture Collection), USA) was cultured with MEM (HyClone) complete medium containing 10% (v/v) fetal bovine serum (Pan), 1% (v/v) penicillin-streptomycin solution (100 U/mL of penicillin G sodium, 100 U/mL of streptomycin) (HyClone),1% (v/v) MEM non-essential amino acid (Gibco). Human monocytic U937 cells (ATCC, USA) were cultured in RPMI 1640 medium (Hyclone) supplemented with 10% heat-inactivated fetal bovine serum (FBS, Gibco, Invitrogen)) and 1% antibiotics (100 U/mL of penicillin G sodium, 100 U/mL of streptomycin), at 37□ and 5% CO_2_ in the incubator. Human monocytes were isolated from PBMC using a magnetic-based monocyte isolation kit (Miltenyi Biotec) and differentiated into macrophages with GMCSF (50 ng/mL; R&D) for 3 d and then RPMI complete medium for 4 d. The OSCC primary cells were obtained from the primary tissues from the Affiliated Hospital of Dalian Medical University. The Ethics Committee approved the collection and use of human samples of Dalian Medical University. Cells were cultured at a laminar flow hood (37□ and 5% CO_2_). The fetal bovine serum needs to be ultra-centrifuged at 100,000 x *g* at 4 ºC for 4 h before removing serum exosomes’ interference in single-cell EV secretion detection.

### Device fabrication

The procedures to fabricate the NOMA microchip are as follows: 1) Fabrication of the hard molds for microwells by patterning SU8-3035 (Microchem) on silicon wafers with photolithography. 2) Spin coating PDMS (base: curing agent = 20:1) on the 30 μm-microwell molds by 1700 rpm to obtain perforated PDMS microwell film. Molding of PDMS microwell (200 μm diameter) with PDMS (5:1 ratio). 3) Align the PDMS microwell (200 μm diameter) slab with perforated PDMS microwell film and bake the assembly at 80 □ for 2 h to realize the irreversible bonding. After curing, the PDMS microchip was peeled off the mold, cleaned for further use.

### Antibody immobilization on the poly-L-lysine coated glass slide

100 μL antibody solution (10 μg/mL) was added to the 1-mm-thick poly-L-lysine coated glass slide. Press the cover glass from left to right to distribute the antibody solution evenly on the glass and incubate it at 4□ overnight. Disassemble the cover glass from the poly-L-lysine glass and block the glass slide with 1% BSA for one h at room temperature. After blocking, the glass slide was washed with DPBS, 50/50 DPBS/DI water, and DI water sequentially. Finally, dry the antibody glass slide with a slide centrifuge and store the antibody-coated slide at 4□ before use.

### Single-cell detection procedures

After cell suspension was collected, they were stained with viability dye Calcium-AM before adjusting the cell density to 1*10^5^ cells/mL. The PDMS NOMA microchip was treated with oxygen plasma for 1 min before use to render the surface hydrophilic. After plasma treatment, 1 mL cell suspension was added onto the NOMA microchip to let the cells settle down to the bottom of the microwell. A fresh cell culture medium was used for flushing excess cells from the surface of the chip. Then the antibody-coated glass slide and NOMA microchip were clamped together and incubated at a 5% CO_2_ incubator at 37□. After incubation, the glass slide captured with secreted protein/EV was removed from the clamp for detection. For time-resolved single-cell secretion analysis, a new antibody-coated glass slide was imposed for another round of detection. The retrieved glass slide was incubated with a cocktail of detection antibodies for 1 hour and stained with streptavidin-APC (eBioscience, 1:100 dilution) afterward for another 30 min. Then it was washed thoroughly with DPBS, 50/50 DPBS/DI water, DI water sequentially. After drying with a slide centrifuge, it was scanned with a GenePix 4300A fluorescence scanner (Molecular Devices) for fluorescence readout.

### Data analysis

The Nikon Eclipse TiE microscope captured images containing cell numbers/positions across NOMA microchips. The resulting secretion fluorescence signals were scanned by GenePix 4300A and analyzed with GenePix Pro software to add and align the microwells array template, followed by the extraction of fluorescence intensity results. The cell counts in the NOMA microchip can be matched with the extracted fluorescence intensity. The mean intensity plus 2×SD of zero cells was defined as the threshold to determine the positive secretion from single cells. Excel (Microsoft), Origin, or Graphpad Prism were used for data analysis.

## Results and discussion

### Design of the high-density NOMA microchip

Microwell is one of the most widely used microchip platforms for single-cell analysis. During the single-cell operation, cell suspension can be pipetted directly on the top of a microwell microchip to trap single cells inside the microwells, which was pretreated with oxygen plasma to be hydrophilic to facilitate cell loading and minimize nonspecific adsorption. The cells would fall into microwells, and the cell distribution follows Poisson statistics. Different types of detection methods can be employed to profile the single cells retained in the microwells. For secretion detection, an antibody-coated glass slide could capture the secretion information from these single cells. Briefly, the antibody patterned glass slide was assembled on top of the microwell microchip for incubation, during which the targeted secretome would be captured. The glass slide would then be removed to finish one round of capture&detection by immuno-sandwich assay. A new antibody-coated glass slide can be imposed onto the microchip for another round of detection to enable sequential measurements from the same single cells. However, due to the disturbance of this glass changing step, ~43% of the single cells in the open microwells would be lost in this process, even with adherent cells(*38*). While for suspension cells, it would be much more challenging to keep single cells in the same microwells during the glass slide changing step when adherent force is absent between the cells and PDMS substrate.

To minimize cell loss during repetitive detection procedures, we innovatively designed the narrow-opening microwell array (NOMA) to solve the significant cell loss problem. **Figure 1A** shows the image of a PDMS NOMA microchip and its enlarged view, which is a two-layered polydimethylsiloxane (PDMS) microwell array. The bottom layer is a microwell array, and the top layer is a corresponding perforated microwell array with a much smaller diameter. Due to the small opening and increased depth of PDMS microwells on the top, the Laplace pressure of the liquid inside the microwell would increase significantly, which would help to confine the single cells trapped inside the microwells. The top PDMS layer, fabricated with spin coating, was characterized with SEM (**Figure 1B & Figure S1**) to ensure that microwells were fully perforated. The top and bottom layers were cured with PDMS at different crosslinking ratios (10:1 and 5:1) and bonded together irreversibly after alignment (**Figure 1C**). The resulting volume of each narrow-opening microwell is ~3 nL (corresponds to 3.3×10^5^ cells/mL cell density for single-cell occupancy), which is sufficient to accommodate single cells inside and keep them in the viable state over days (**Figure 1D**). The total number of microwells is 4000, ensuring high throughput analysis.

**Figure 1.**
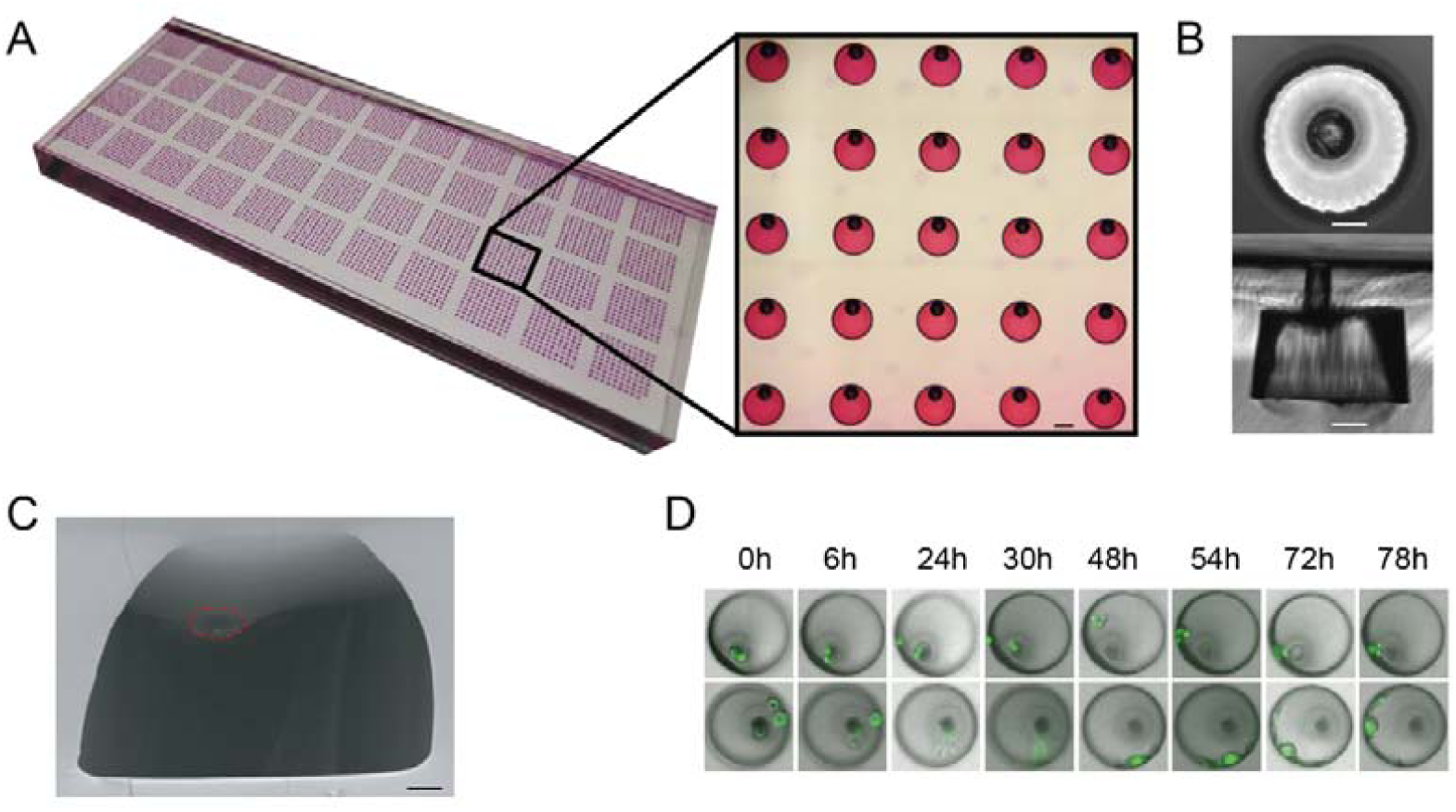
Design of the NOMA microchip. A. Photograph of a NOMA microchip and its enlarged view (filled with food dye to facilitate visualization). B. Optical micrograph showing the enlarged view of the individual microwell with a narrow opening, including the top and the cross-section views. The scale bar is 50 µm. C. SEM characterization of the cross-section view of the microwell with a narrow opening, in which the perforated microwell was circled. D. SCC 25 cancer cells (stained with membrane dye DiO) inside NOMA microchip showed excellent viability over days.

### NOMA microchip realizes robust, high-efficiency retention of single-cells

To characterize the NOMA microchip performance in single-cell retention, we designed the NOMA microchips with different diameters of narrow top opening, including 20, 30, 50, 60, 80, and 100 µm (**Figure 2A**), and the bottom microwell is 200 µm in diameter. The results showed that the single-cell occupancy was around 21%, which is almost the same when the narrow opening increased fivefold from 20 µm to 100 µm. We chose 30 µm as the diameter of the narrow top opening for the following experiments to minimize the microwell volume for sensitive detection and facilitate cell handling. We then evaluated the cell retention efficiency by changing glass slides on the top of the NOMA microchip repetitively. We found that ~98% of adherent SCC25 tumor cells (**Figure 2B**) would be kept in the same microwells after the glass changing step, which is significantly higher than previous reports (p<0.0005, **Figure S2**). Two different researchers verified the retention results independently, which showed no significant difference between each other (98% vs. 99%, p=0.39), demonstrating the method’s robustness. Notably, the excellent cell retention efficiency works not only for adherent cells but also for suspension cells, verified with U937 suspension cells (average diameter: ~10 μm). We found around 97% of the cells would be kept in the same microwells after the glass changing step (**Figure 2C**), and the result was also validated by two researchers independently (97% vs. 97%, p=0.67), further demonstrating the robust, excellent single cell retention efficiency of NOMA platform. A hypothetical comparison of the single-cell retention between the open microwell and the NOMA platform reveals that only 3% of single cells would be kept inside the open microwells after six rounds of repetitive detection (**Figure 2D**). In comparison, 89% of single cells could still be retained in the NOMA microchip, demonstrating a significant improvement in single-cell retention efficiency.

**Figure 2.**
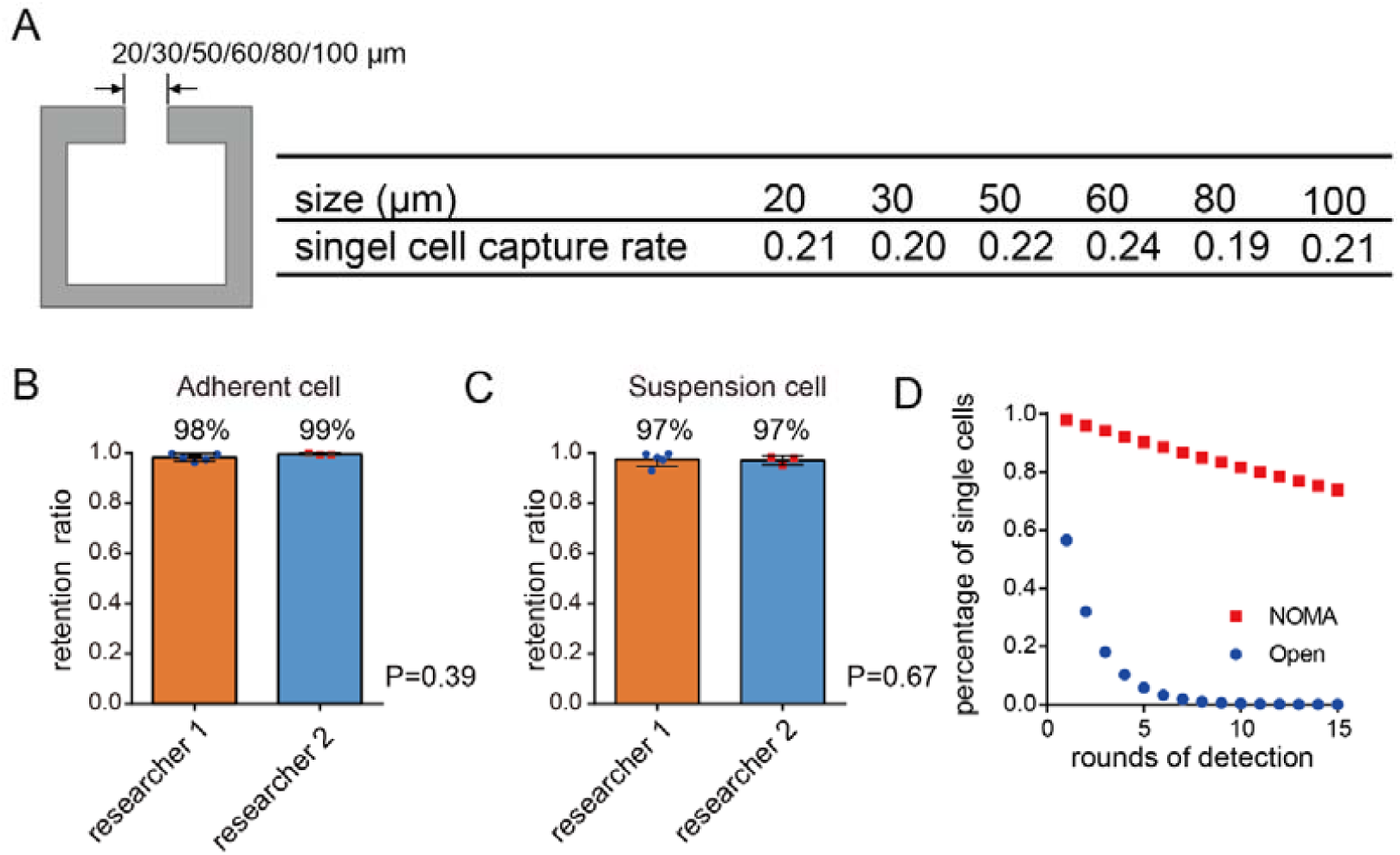
Characterization of the NOMA microchip. A. Evaluation of NOMA microchips with different diameters of the narrow top opening for single-cell capture rate. B-C. Both adherent cell (B) and suspension cell (C) showed excellent single-cell retention efficiency during repetitive glass slide changing procedures, and the results were verified by two researchers independently. D. Hypothetical comparison between NOMA microchip and open microwell microchip in cell retention performance after repetitive detection procedures based on their respective cell retention efficiency.

### Measuring the secretion sequentially from the same single cells reveals the dominance of the digital secretion pattern

Profiling cell secretion profiling, including cytokines, chemokines, etc., provides a direct way to evaluate the functional cell states for different cells, including immune cells, cancer cells(*39–41*). Due to the dynamic characteristic of protein secretion, secretion detection from the same single cells at different time points is essential, which would provide unique information, besides the magnitude and intensity, to reveal the new heterogeneity in persistence underlying secretion. We applied the NOMA platform for tracking the secretions from the same single-cells sequentially to show the time-resolved heterogeneity in secretion.

We firstly used macrophage cells in response to LPS stimulation as a model system to profile the protein secretion sequentially from the same single cells. Primary human monocytes from a healthy donor were isolated from PBMC with magnetic sorting and further differentiated into macrophages with GMCSF(*18, 39*). Macrophage cells were challenged with 100 ng/mL LPS before the single-cell assay, which would activate the pathogen recognition by binding to the TLR4 receptor and stimulate the innate immune response against Gram-negative bacteria(*17*). An antibody-coated glass slide was imposed on the PDMS microwells to profile protein secretion, fabricated by immobilizing antibodies associated with innate immune responses, including IL-8, IL-6, onto poly-L-lysine coated glass slides under the static condition at room temperature. The antibody pairs used were titrated with corresponding recombinant proteins (**Figure S3**&**Table S1**). After measuring protein secretion from single macrophage cells for 4 hrs, we removed the antibody glass slide and then put another glass slide coated with the same panel of antibodies (or a different panel of antibodies) to measure protein secretion from the same single cells for another 4 hrs. We successfully obtained secretion data from 903 single cells at two-time points along with the LPS stimulation, which could be visualized with line graphs showing the oscillatory change of protein secretions from the same single cells (**Figure 3A**).

**Figure 3.**
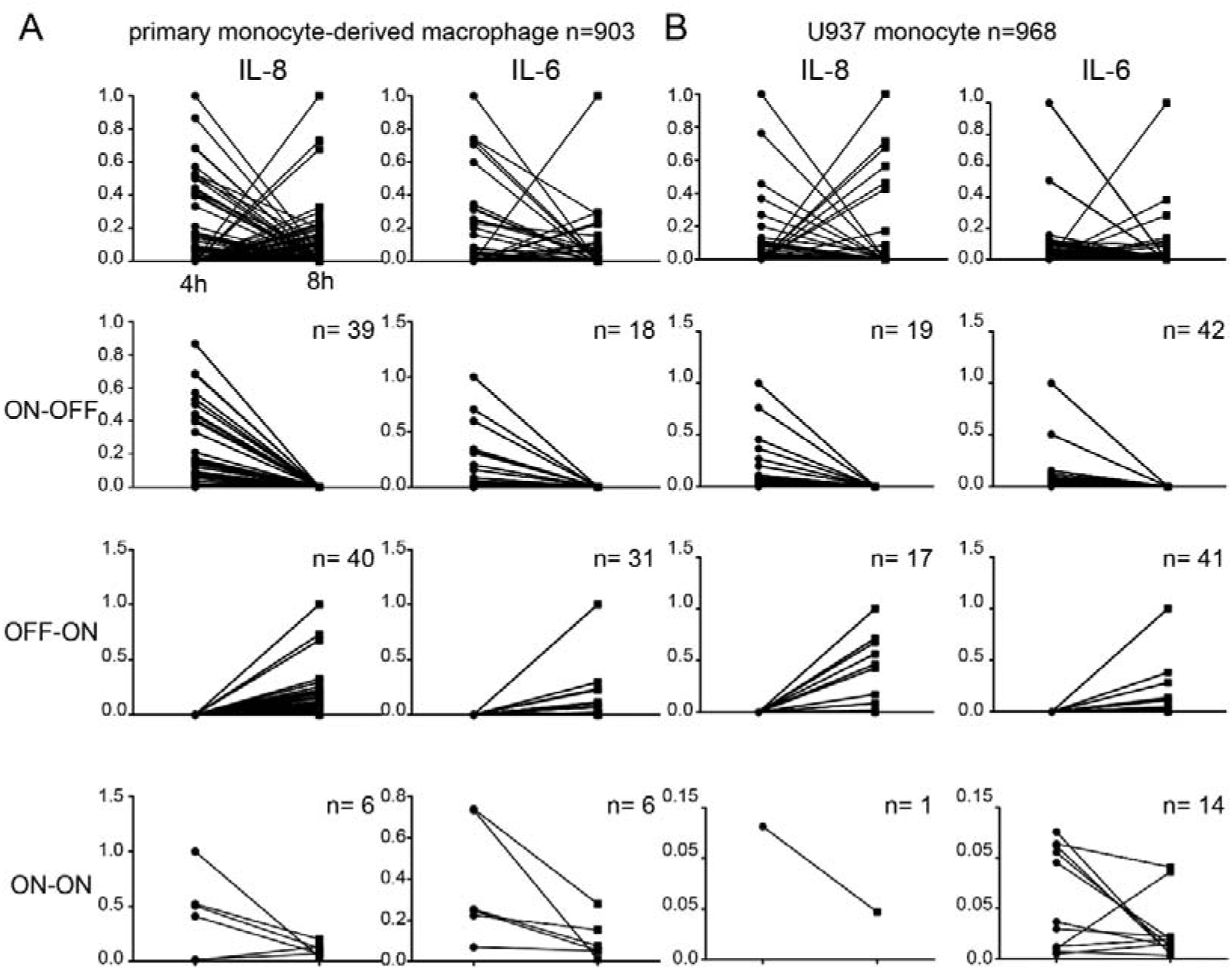
Line graphs showing the sequential readout of protein secretion information (IL-8, IL-6) from the same single cells (primary monocyte-derived macrophage(A) and U937 monocyte) revealed that protein secretion from single cells could be classified into three different patterns based on their change at two different time points: on-off, off-on, and all on. The y-axis is normalized based on the maximum value of each secretion.

Interestingly, we found all single cells positive with protein secretion at either of two time points could be classified into three secretion patterns based on their dynamic change: ON-OFF, OFF-ON, and ON-ON. For example, we observed that 39 single cells for IL-8 secretion during the first 4 hrs would be negative for IL-8 secretion in the second 4 hrs. Similarly, 40 cells positive for IL-8 secretion in the second 4 hrs were negative for IL-8 secretion in the first 4 hrs. Only six cells were positive in IL-8 secretion at both the first 4 hrs and the second 4 hrs. Similar secretion behaviors were also observed from IL-6 protein. These data suggest that individual cells secrete proteins for relatively short periods, and protein secretions are dominated by digital secretion patterns (ON-OFF & OFF-ON, 93% for IL-8, 89% for IL-6). The results are consistent with a previous study that revealed the digital activation of transcription factor nuclear factor NF-kB in response to activation at the single-cell level(*42*). Here we used the NOMA platform to demonstrate the stochastic oscillation pattern within single cells at the functional protein level, complementing the observation at the transcriptional level.

We also applied the NOMA microchip to profile the protein secretions from suspension cell U937 (**Figure 3B**) and OSCC cancer cells (**Figure S4**) to assess the broad applicability of the platform. We found the secretion patterns in these two types of cells could also be resolved into ON-OFF, OFF-ON, and ON-ON patterns, in which digital secretion patterns (ON-OFF& OFF-ON, U937 monocyte: 97% for IL-8, 86% for IL-6; OSCC cancer cell: 82% for IL-8, 85% for IL-1b) dominated the secretions. Collectively, these results revealed the existence of three different secretion patterns among different types of cells, indicating the intrinsic dynamic feature of cell heterogeneity in protein secretion. These observations also revealed the dominant roles of digital secretion patterns in these cells, which are difficult to observe with single-cell analysis technologies based on a single snapshot measurement, demonstrating the unique merit of the sequential measurements from the same single cells with NOMA microchip.

### Longitudinally tracking of the single-cell integrative secretions over days

The unique merit of the NOMA platform is its high fidelity in single-cell retaining capability, making it possible to track the information from the same single over a long time, which is challenging to realize with existing technologies. To demonstrate, we used the platform for longitudinally tracking of the single-cell integrative secretion over days. Oral squamous cell carcinoma cell line (SCC25) was used as the model cell line, which would secrete cytokines or EVs continuously. After SCC25 cells were enclosed as single cells into microwells, the antibody-coated glass side was used to capture the secretions from these single-cells twice each day (4 hours incubation for each detection window), three days in total (**Figure S5**). To obtain integrative secretion information from the same single cells, we immobilized IL-8 antibody for cytokine capture and CD81 for EV capture simultaneously on the same glass slide. Titration tests with EV standards were completed to characterize detection sensitivity (**Figure S6**). We successfully obtained IL-8 and ^CD81+^EV secretion data from 377 single-cells across six different time points, summarized with line graphs (**Figure 4A-B**) or other analysis tools (**Figure S7**) to show the changes of secretion from each cell. Around 35% of cells released IL-8 secretion more than once during the detection time, and most of the IL-8 secretion follows the digital pattern, which is consistent with our previous observation. Most of the single cells would only produce IL-8 protein during a 1-time window. A small population of cells (~2%) was quite active to secrete IL-8 protein more than three times, indicating the presence of “super secretors” within the cell population that could secrete proteins more persisting for an extended period. Similarly, we found 18% of cells produced ^CD81+^EV secretion more than once during the detection, and the digital pattern also dominated EV secretion. Only two cells (0.5%) were able to secrete ^CD81+^EV more than two times, further confirming the presence of persisting active secreting cells in the population. These results reveal the dynamic heterogeneity underlying cell secretion that a few cells could secrete cytokines or EVs multiple times, which could not be observed with snapshot measurements.

**Figure 4.**
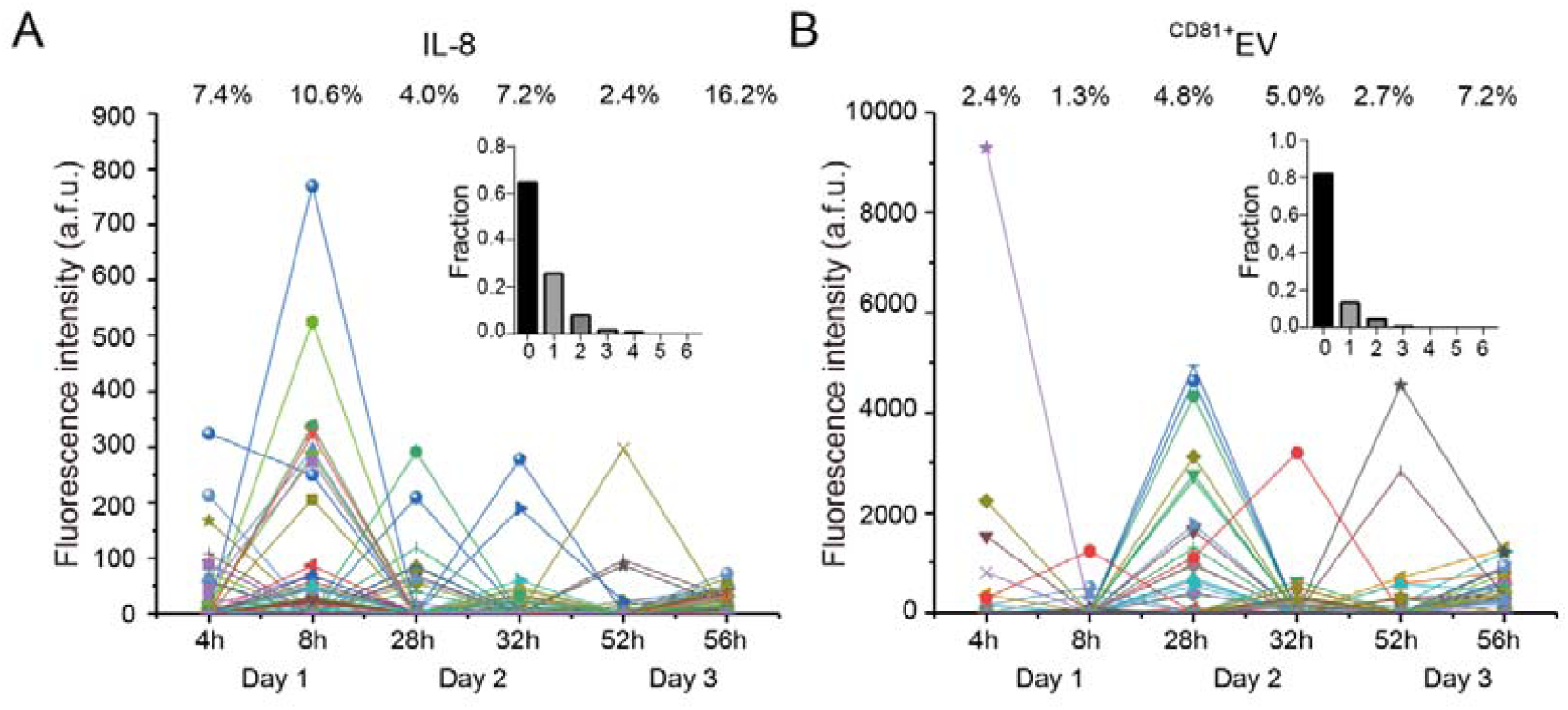
Longitudinally tracking of integrative secretions from single cells over days. Line graphs showing the change of individual secretions (A: IL-8; B: ^CD81+^EV) from 377 single cells over three days.

Previous studies have revealed polyfunctional immune cells within the cell population are critical players in protective immunity and immunotherapy, demonstrating the significance of secretion heterogeneity. However, it is still difficult to evaluate the persistence of secretion at the same single-cell level. Besides, how the heterogeneity in the continuation of secretion would affect the cell function is yet to be explored. However, these studies are mainly hindered by the lack of research tools. The NOMA microchip that could realize the robust, high-efficiency single-cell retention demonstrated itself potentially be a valuable tool to widely used for deep phenotyping dynamic heterogeneity and persistence of secretion. Besides, it is also possible to recover the single-cells of interest selectively from the NOMA microchip (**Figure S8**) and be further coupled with other single-cell manipulation or detection methods, such as genome editing or RNA-seq, etc., to obtain more information from the same single cells.

## Conclusion

We reported a NOMA microchip to track the information from the same single cells, which realized the retention of over 97% of single cells with both adherent cells and suspension cells during repetitive detection procedures. The unique merit of this single-cell microdevice is the robust, high-efficiency single-cell retention to keep single cells alive in the dedicated spatial locations after rounds of detection. With this platform, we realized the sequential measurements of the same live single-cell to realize the tracking of single-cell secretion longitudinally, revealing three different secretion patterns and the dominant roles of digital secretion patterns among different types of cells. The highly robust cell retention capability also makes it possible to recover the single-cells of interest selectively based on their secretion profiles and be coupled with other single-cell manipulation or detection methods for additional studies in the future. The number of secretome parameter can also be further increased by the combination of barcoding technology, including antibody barcode and bead barcode. This platform represents a novel and powerful tool broadly applicable for different cell types, including immune cells (B cell, T cell), tumor cells (circulating tumor cells), for informative monitoring of the same single cells in basic and clinical research.

## Supporting Information

Supporting Information is available and includes platform characterizations and single-cell secretion analysis with tumor cells.

## Conflict of Interest

There is no conflict of interest to report.

## Funding Information

This research was made possible as a result of a generous grant from the National Natural Science Foundation of China (Grant No. 21605143, 21874133, 31927802, 82073001), Youth Innovation Promotion Association CAS (Grant No. 2018217), and funds from the Dalian Institute of Chemical Physics (Grant No. DICP I201908).

## Acknowledgment

The authors wish to acknowledge Professor Chaoqun Rao for his helpful discussions regarding the experimental design.

## Supporting information

### Supporting figures

**Figure S1.**
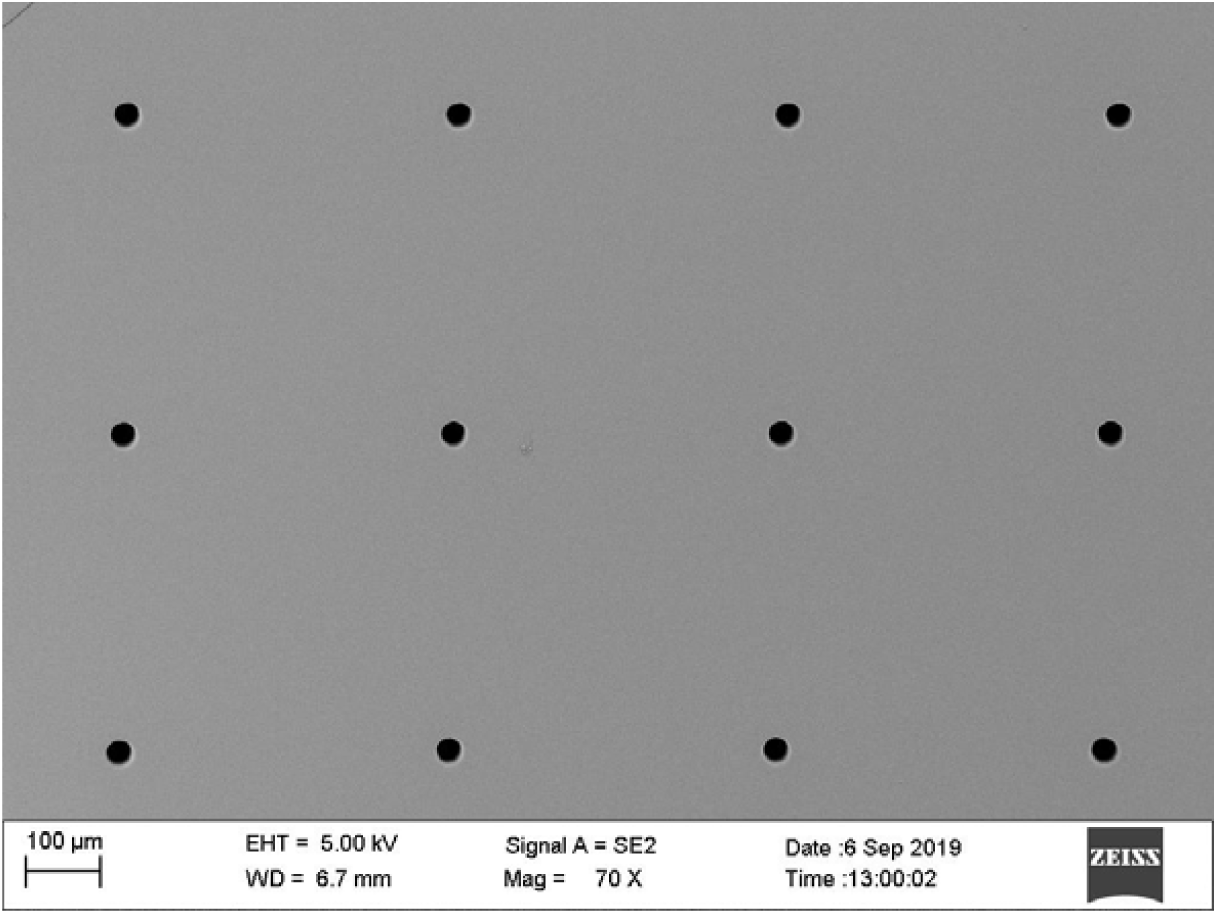
Characterization of perforated PDMS microwells (diameter: 30 µm) with SEM.

**Figure S2.**
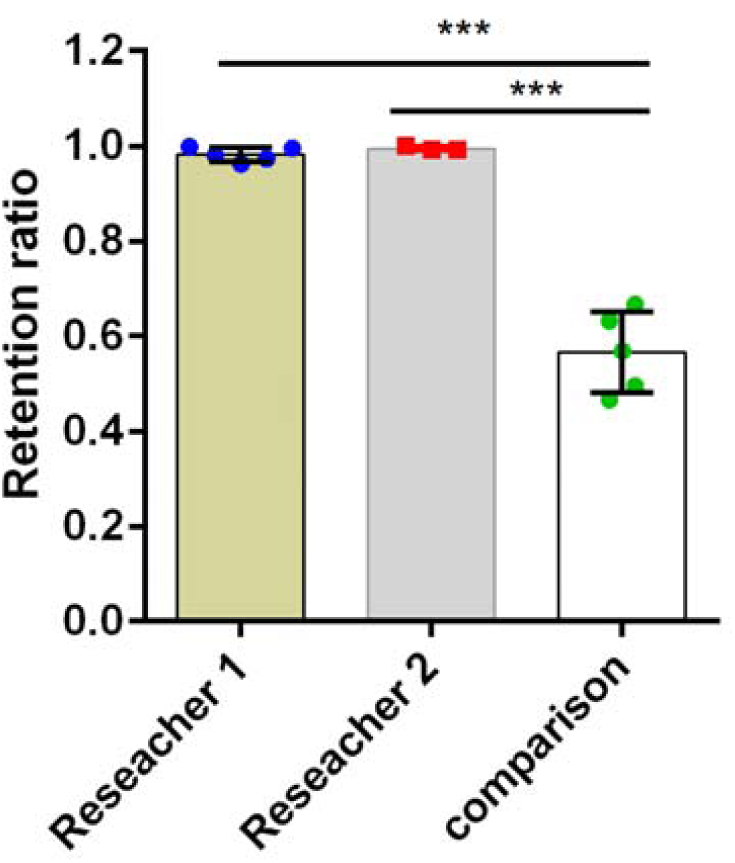
Comparison of cell retention efficiency between NOMA microchip and open-microwell based microchip during repetitive glass changing steps.

**Figure S3.**
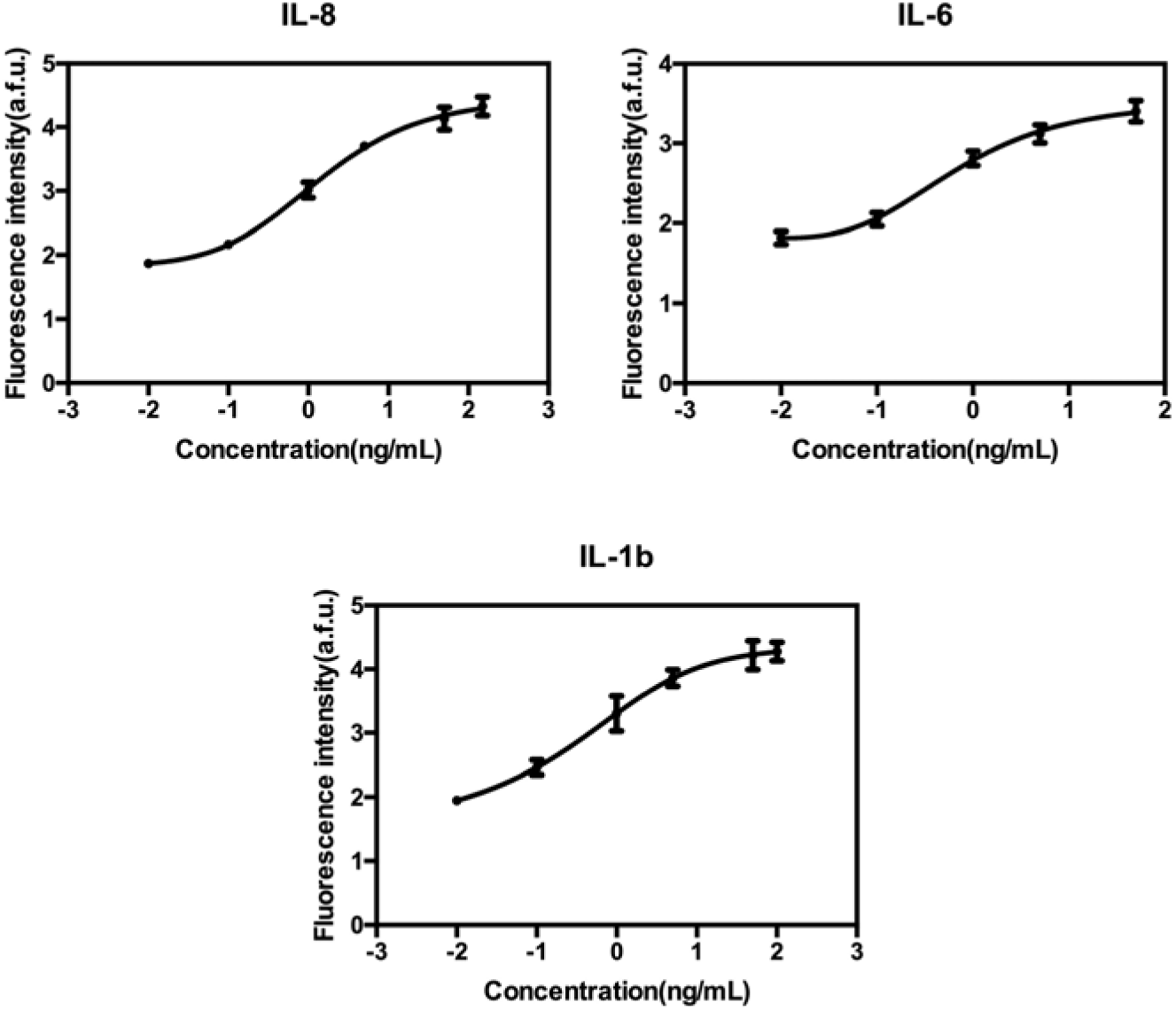
Calibration curves were obtained with corresponding recombinant proteins (x, y axes: log-scaled, the intensities were averaged from 6 spots for each data).

**Figure S4.**
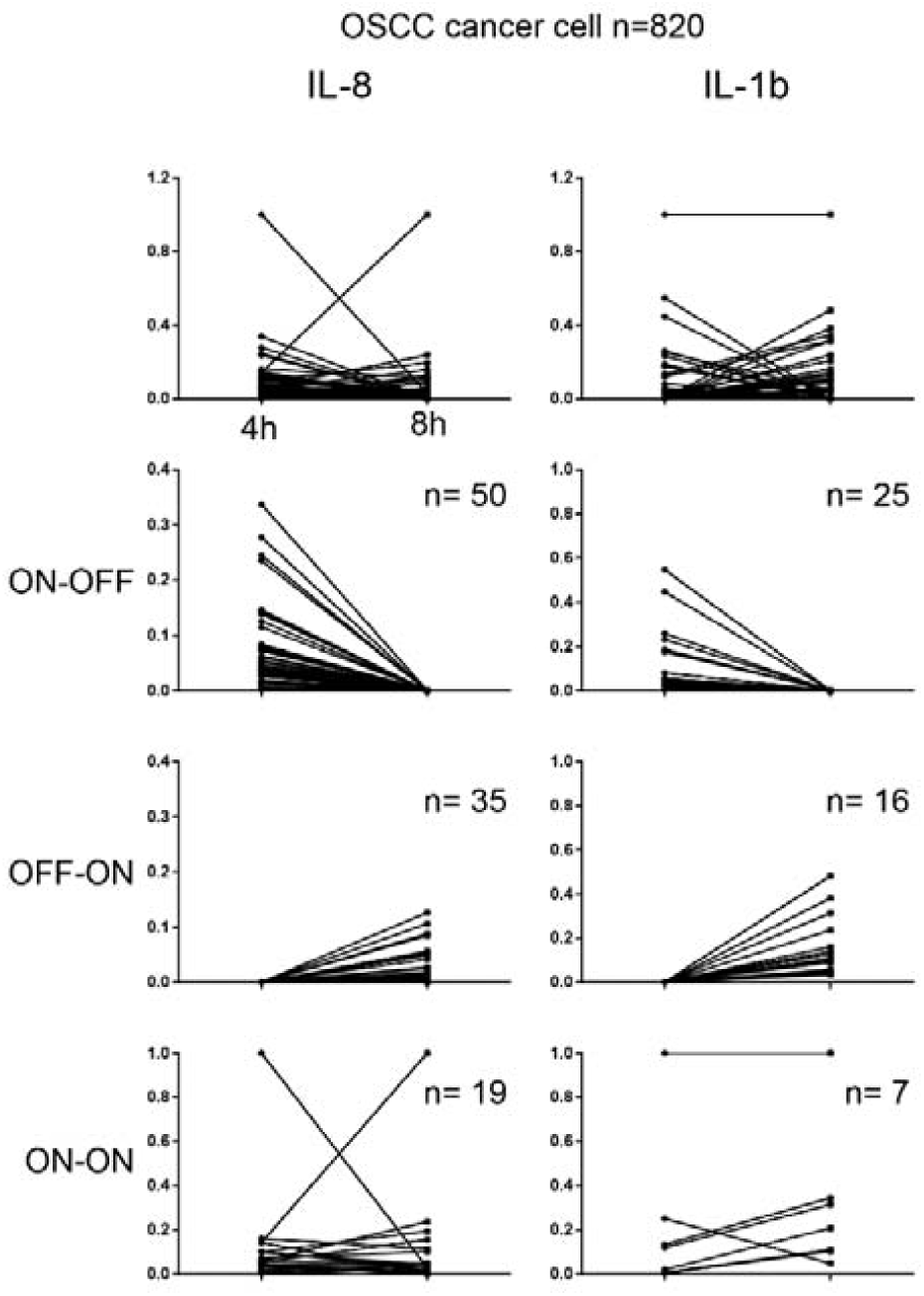
Sequential measurements of protein secretions(IL-8, IL-1b) from the same OSCC single cells (n=820) revealed that protein secretions from single cancer cells could also be classified into three secretion patterns based on their change: on-off, off-on, and all on, and the digital secretion patterns (on-off, off-on) dominated the secretion behaviors.

**Figure S5.**
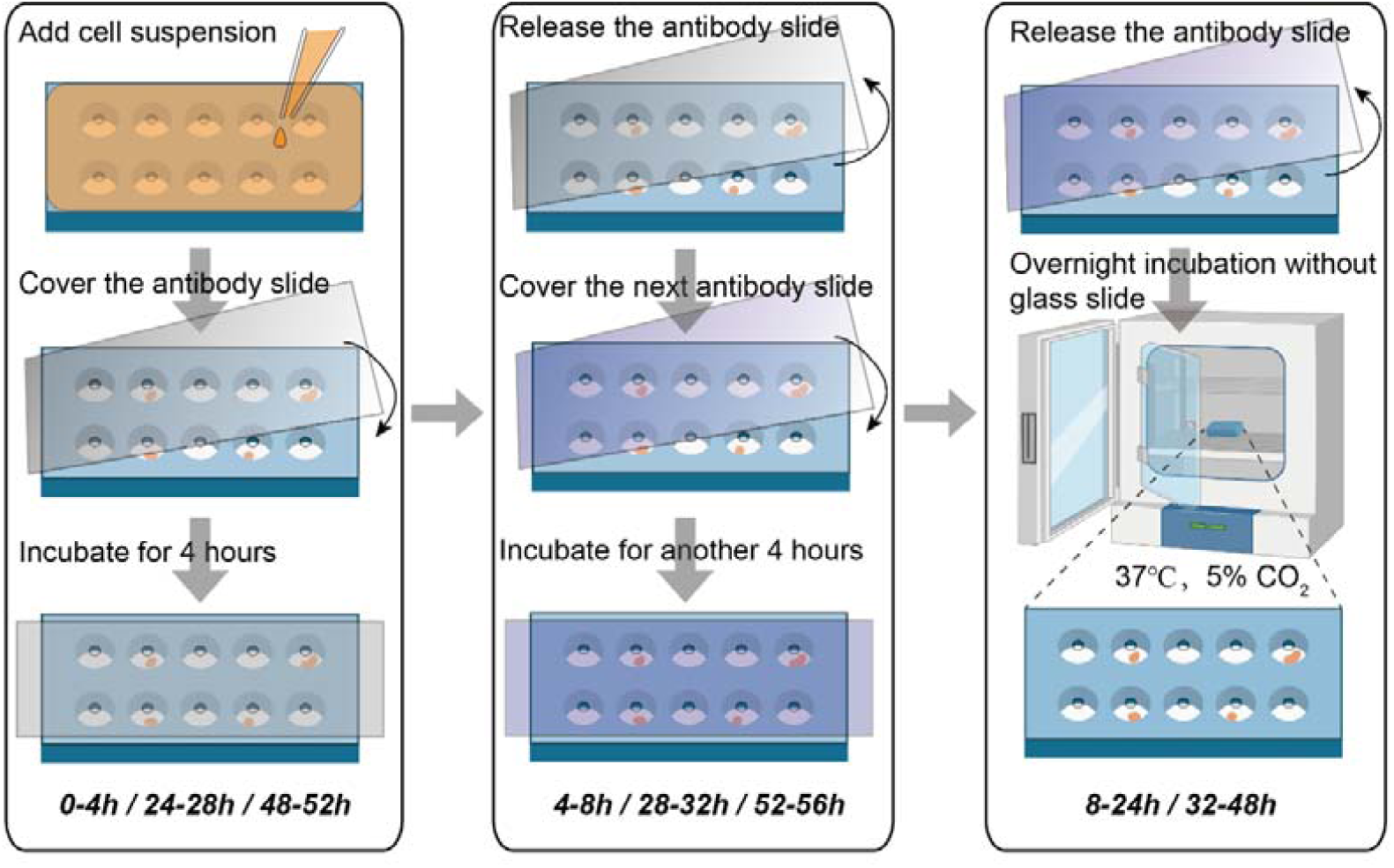
Illustration of the detection procedures to follow the secretions from the same single cells at different time points over three days.

**Figure S6.**
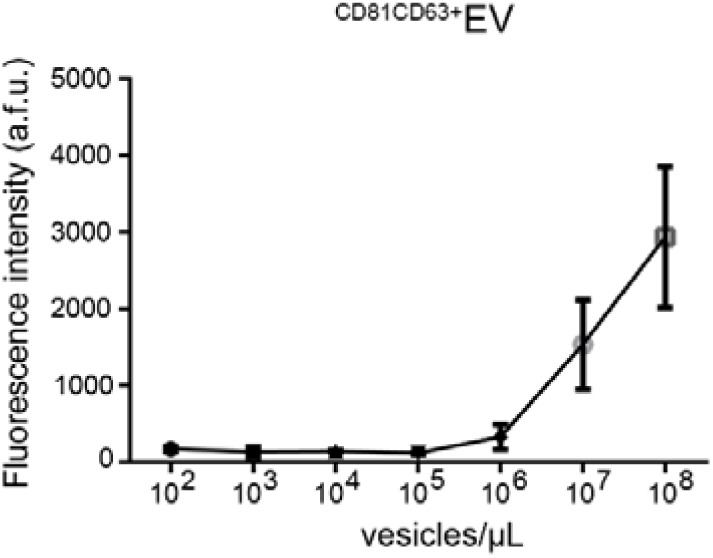
Sensitivity characterization of the antibody-based EV detection with EV standard (ExoStd™ Lyophilized Exosome Standard, Biovision, USA) reveals that the lowest detection concentration for EV is ~10^5^ EVs/μL. Considering the volume of an individual microwell in the NOMA microchip is ~3 nL, the lowest detectable number of EVs is ~300 vesicles/microwell.

**Figure S7.**
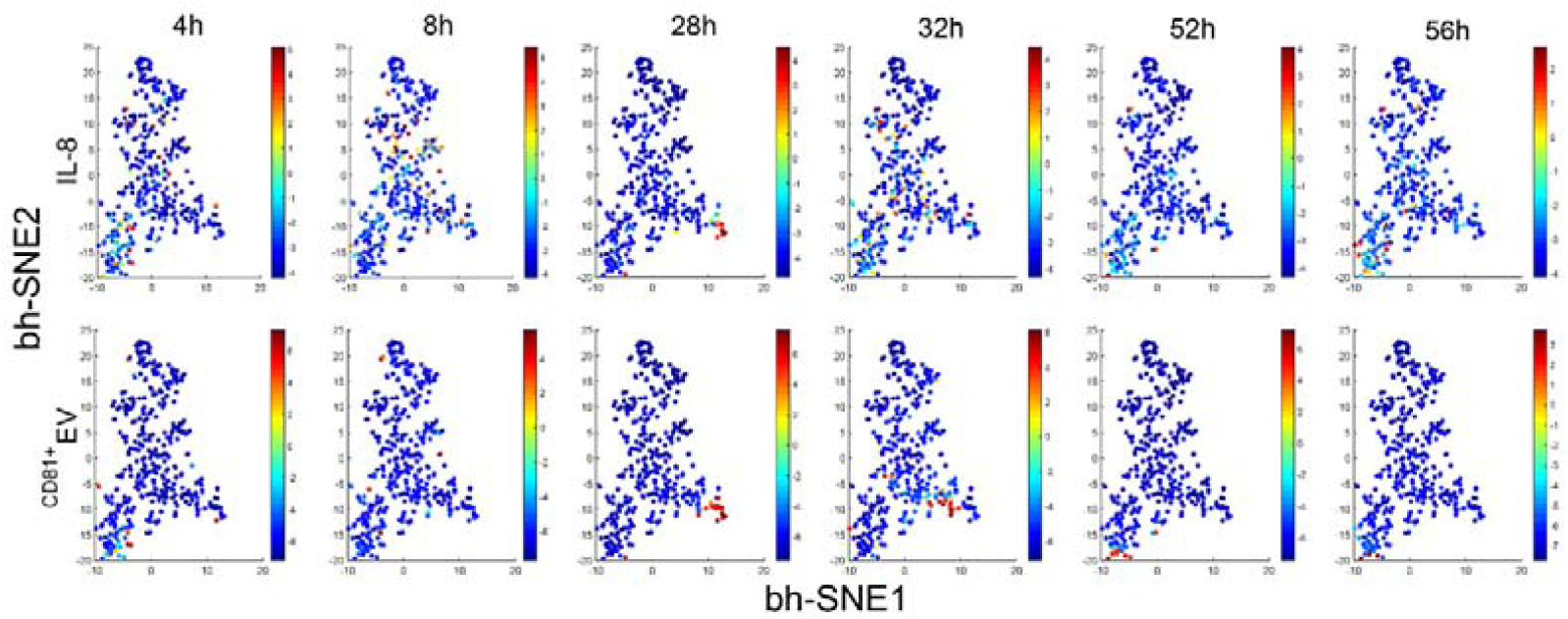
Visualized clustering analysis with viSNE to reveal the protein and EV secretions from the same single cells at different time points. All the single-cell data at six different time points with both protein secretion and EV secretion were combined into a unique dynamic single-cell data metric to be visualized with viSNE, which is a high-dimensional data analysis tool based on the t-Distributed Stochastic Neighbor Embedding algorithm. Interestingly, we found EV secretion and protein secretion are from into different cells, except for 28 h, during which they were nicely correlated with each other at this time point (Pearson r=0.89). This was not observed previously, highlighting the value of a dynamic detection platform with the single-cell resolution to reveal how biological systems respond and evolve.

**Figure S8.**
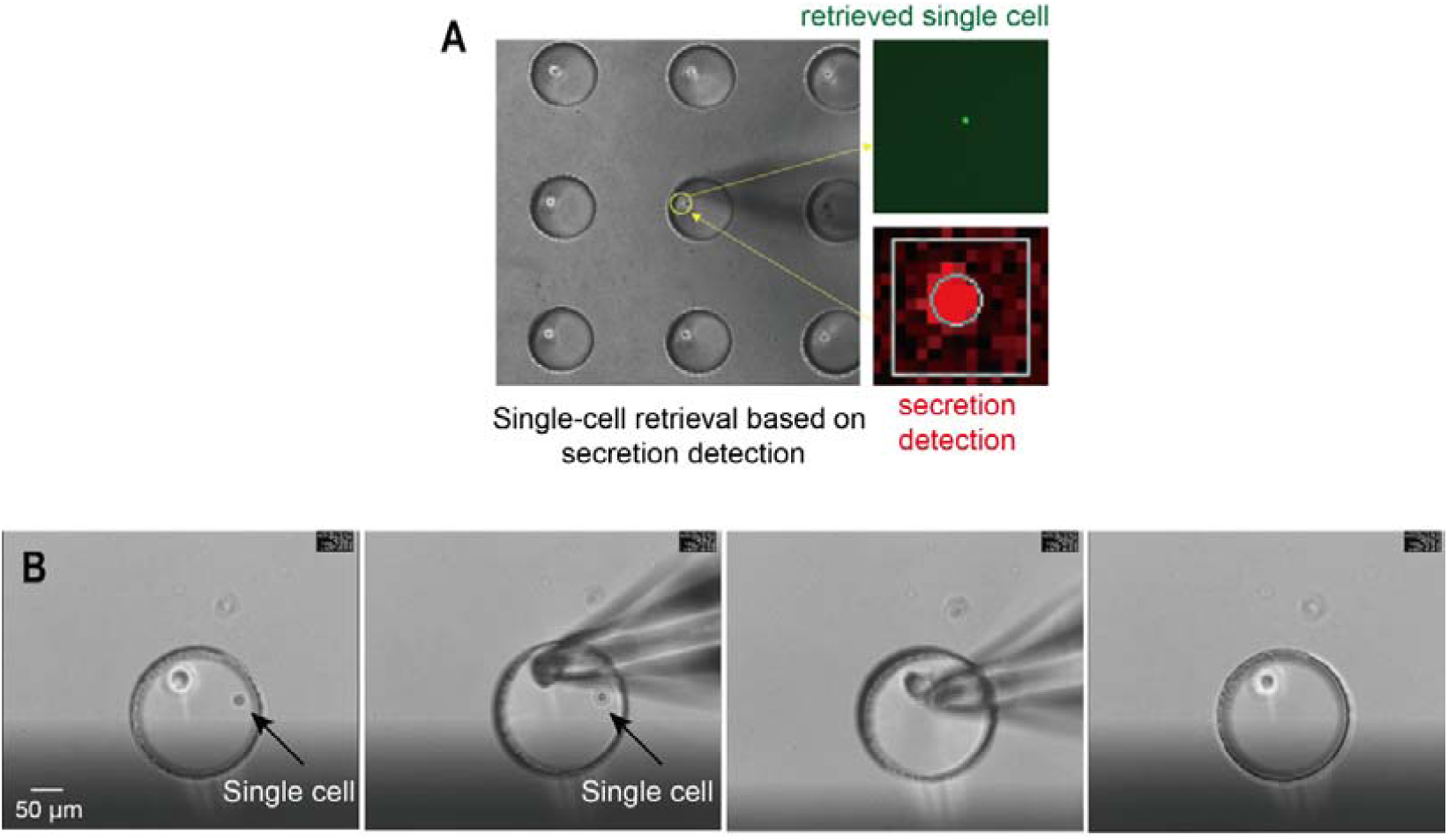
Single-cell retrieval from an individual microwell in the NOMA microchip following secretion detection. A. A representative retrieved single cell. B. Video clips showing the retrieval process.

**Table S1.**
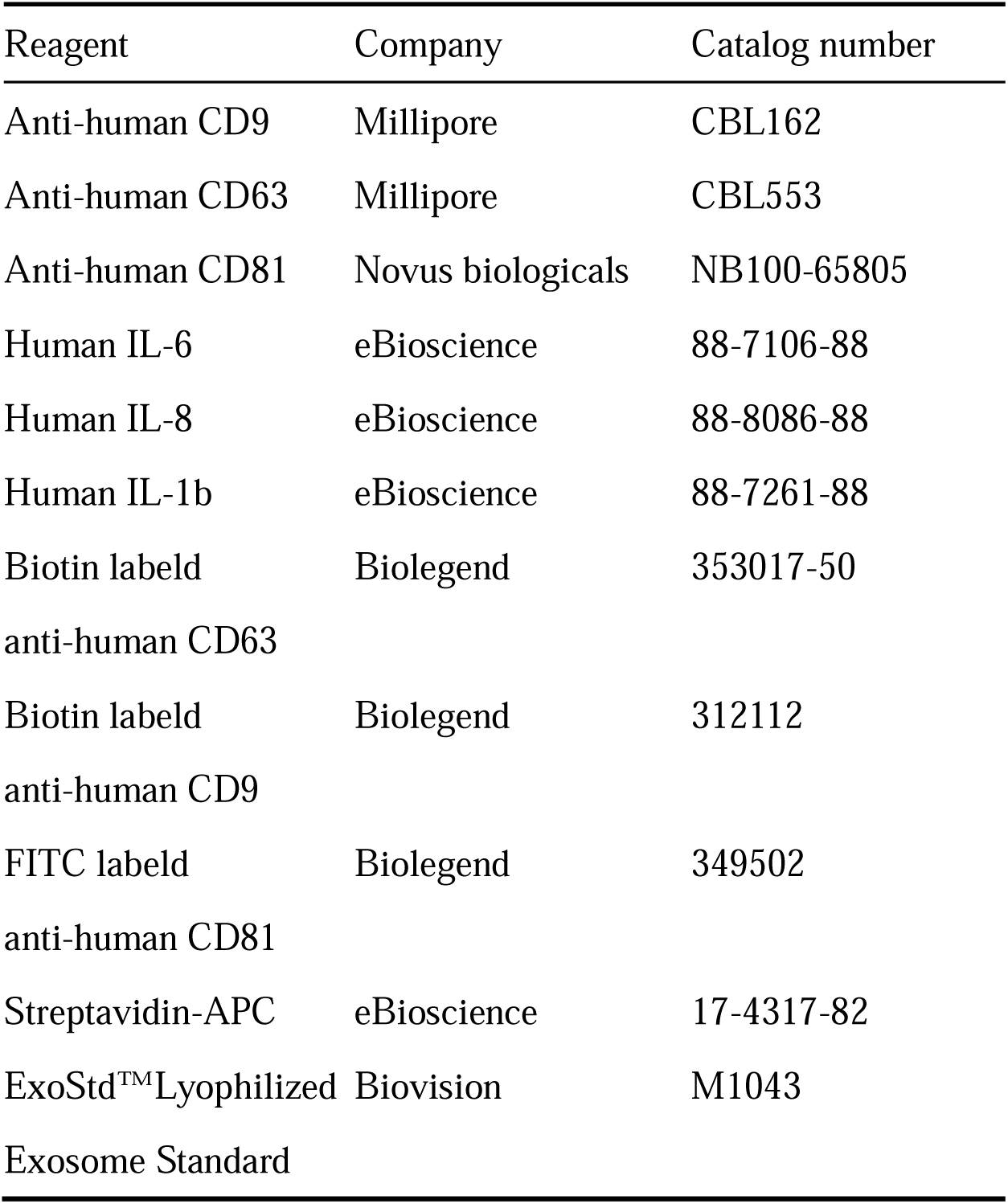
Summary of key reagents used in this study.

## Notes

### Competing Interest Statement

The authors have declared no competing interest.

